# Viscoelastic dissipation stabilizes cell shape changes during tissue morphogenesis

**DOI:** 10.1101/107557

**Authors:** R Clément, C. Collinet, B. Dehapiot, T. Lecuit, P.-F. Lenne

## Abstract

Tissue morphogenesis relies on the production of active cellular forces. Understanding how such forces are mechanically converted into cell shape changes is essential to our understanding of morphogenesis. Here we use Myosin II pulsatile activity during *Drosophila* embryogenesis to study how transient forces generate irreversible cell shape changes. Analyzing the dynamics of junction shortening and elongation resulting from Myosin II pulses, we find that long pulses yield less reversible deformations, typically a signature of dissipative mechanics. This is consistent with a simple viscoelastic description, which we use to model individual shortening and elongation events. The model predicts that dissipation typically occurs on the minute timescale, a timescale commensurate with that of force generation by Myosin II pulses. We test this estimate by applying time-controlled forces on junctions with optical tweezers. Our results argue that active junctional deformation is stabilized by dissipation. Hence, tissue morphogenesis requires coordination between force generation and dissipation.

---

The course of animal development is a succession of morphogenetic movements, which require the activity of force-generating cortical components that exert mechanical forces at the cellular scale (1,2). Classic examples include cell intercalation, in which polarized activity of Myosin II (MyoII) motors drives tissue elongation (3,4), or apical constriction in which MyoII recruitment drives tissue folding and invagination (5). Thus, MyoII contractility causes cells and tissues to undergo massive deformation. While global tissue movements last one to a few tens of minutes, individual cell deformations are often shorter events that rely on transient forces resulting from pulsatile contractions of the acto-myosin network (5–8). A key question is thus how these deformations are stabilized so as to ensure persistent deformations. This requires a mechanical understanding of how cells and tissues escape elastic recoil once contractile stresses (e.g. forces) are no longer applied. Indeed elastic materials store elastic energy when submitted to deviations from their reference configuration, and recoil towards this configuration upon release. In addition, application of a constant force yields a constant deformation (e.g. deviation from the reference configuration), and going further away from the reference configuration requires ever-increasing forces, which in the case of biological systems might cause fracture or loss of cellular integrity (9). On the other hand, viscous materials do not store elastic energy and thus have no reference configuration. As a result, deformation keeps increasing as long as a force is applied, and no recoil is observed upon release. Such lack of robustness to short-lived or abnormal forces would be deleterious to a biological system. It was noted long ago that the physical nature of cells and their cortex-a dynamic, cross-linked polymer with rapid turnover-should confer viscoelastic, time-scale dependent mechanical properties to embryonic tissues (10–13). Thus, a likely mechanism to achieve irreversible deformation is the gradual dissipation of elastic energy during the morphogenetic process (9). Dissipation causes a drift of the reference configuration, allowing forces exerted transiently to generate persistent shape changes. In order for the latter to be true, dissipation should occur on time scales typically shorter, or commensurate with, but definitely not much longer than the timescale of force generation. Hence it is often assumed that tissues behave as fluids on developmental timescales. Experimentally, the viscoelastic nature of single cells or cell assemblies have been assessed with a variety of techniques (14–17) and recent works using suspended epithelial monolayers (18) confirm that dissipation occurs upon application of a constant deformation, typically on the minute timescale (19). Yet very few studies involve direct mechanical measurements *in vivo* in the context of morphogenesis. In a previous study using the *Drosophila* embryonic epithelium as a model system, we delineated the mechanics of cell contacts at short time scales, typically below a few seconds, showing that the response was mainly elastic and damped by fluid friction in the cytosol (20). However, longer time scales remain largely unexplored, due to the technical challenge of actively probing cell mechanics in live embryos over long periods of time. Thus a mechanical analysis at morphogenetic time scales and an estimate of the relevant dissipation timescales in a morphogenetically active tissue are still lacking.

Here, we take advantage of the irreversible shortening and elongation of cell junctions in response to pulsatile Myosin II activity to estimate the typical timescale of dissipation at cell contacts. Thereby, we sought to understand how transient forces can generate irreversible deformation during this morphogenetic process.

Planar-polarized MyoII activity was shown to drive cell intercalation and participate in tissue extension in the presumptive ectoderm (3,4). Bursts, or pulses, of MyoII activity along junctions aligned with the dorso-ventral axis (Figure 1A) were shown to gradually shorten junctions, acting as a “mechanical ratchet” (8), and eventually resulting in the disappearance of vertical cell contacts (Figure 1B,D,F). Similar bursts, located in adjacent anterior and posterior cells in the vicinity of a junction’s vertices, were then shown to favor the formation and gradual elongation of new junctions along the antero-posterior axis (Figure 1C,E,G) (21). While it is clear that MyoII is necessary to generate the force driving deformation, the mechanism by which these deformations are stabilized remains elusive. Previous studies showed that efficient shortening requires robust activation of MyoII in junctions (8,22,23). In the following, we introduce a minimal viscoelastic model of the so-called mechanical ratchet, which shows that active MyoII-driven deformations can be stabilized by dissipation. Analyzing junctional shortening and elongation in response to pulsatile forces generated by MyoII, we estimate the typical timescale of dissipation, which separates the elastic, reversible regime from the viscous, irreversible regime. We show that it is typically one minute, and confirm this estimate with optical tweezers experiments in which a controlled external force is applied to junctions.

## Results

Our working hypothesis is that cell junctions are viscoelastic with a short-term elastic response and a long term viscous response due to dissipation, allowing deformations generated by transient MyoII activity to be stabilized. Note that cells junctions, lined by a complex network of treadmilling actin filaments crosslinked by a number of turning-over proteins, are likely to undergo dissipation on a distribution of timescales. For the sake of simplicity, we assume in the following that dissipation occurs on a single typical timescale τ.

Junction length dynamics during contractile pulses is correlated with the fluctuation of MyoII intensity (5,8), which is known to generate forces at junctions (24). Therefore we ultimately want the model to predict the associated dynamics of MyoII intensity *M*(*t*) and junction length *l*(*t*). Since a mechanical model relates force to deformation, we first need to relate MyoII intensity to force *f*(*t*). MyoII recruitment at the cortex requires activation and is used as a proxy for motor activity. While it is largely assumed that force increases with MyoII activity, the precise scaling between MyoII and force production is unknown. For the sake of simplicity, we assume in the following that force is proportional to the excess of MyoII (with respect to MyoII constitutive level *M*_0_ observed in absence of pulses), so that *f* = ± *α* (*M* – *M*_0_). In practice we assume that *M*_0_ is the minimum of *M*(*t*), and the ± sign stands for shortening (−) or elongation (+). Finally *α* is an unknown scaling factor. From there, the dynamics of a viscoelastic junction submitted to MyoII-driven shortening or elongation is given by the following reformulation of the Maxwell model of viscoelasticity (see supplementary information for details):

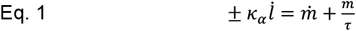

In Eq.1, *κ_α_* is the ratio between the elastic modulus *κ* and the constant *α*, and m = *M* – *M*_0_. Dots denote time derivatives, so that 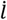 is the rate of elongation, and *ṁ* the rate of MyoII recruitment. Note that here we neglect variations of tension in adjacent junctions, as well as the contribution of viscous drag in the cytosol, which occurs on timescales much shorter than MyoII fluctuations (20). The model has two free parameters: *κ_α_*, which controls how much MyoII can elastically deform a junction, and τ, the typical dissipation timescale. In the case of shortening, *M* is MyoII fluorescent intensity measured in the vicinity of the junction (Figure 1H), where it is supposed to be the most effective to generate a shortening force. In the case of elongation, junctions have very low MyoII levels, and we measure MyoII intensity *M* in adjacent anterior and posterior cells, in the vicinity of vertices (Figure 1I), where it was shown to drive the growth of the new junction (21,25).

**Figure 1.**
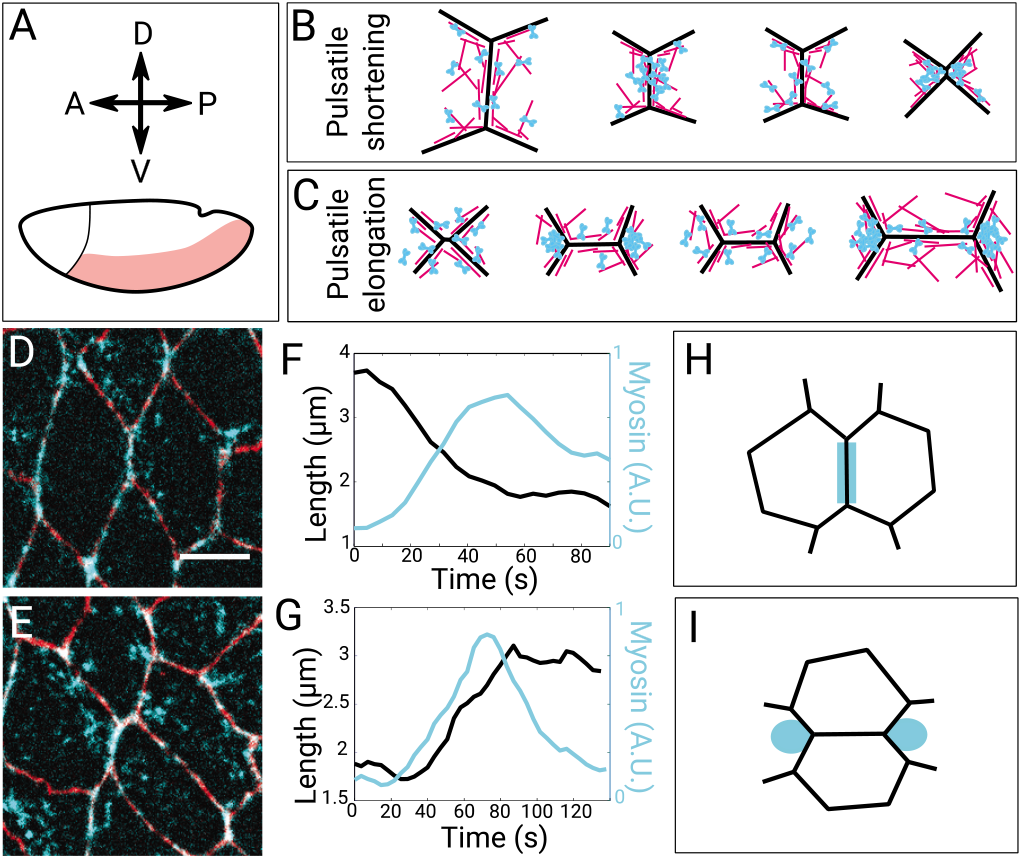
Junction shortening and elongation by MyoII polarized activity. **A.** Schematics of the germband of the *Drosophila* embryo. A, P, D, V symbolize the anterior, posterior, dorsal and ventral directions. **B.** Cartoon of junction shortening driven by MyoII pulsatile activity. Actin is in magenta, MyoII in cyan. **C.** Cartoon of junction elongation driven by MyoII pulsatile activity. Actin is in magenta, MyoII in cyan. **D.** Junction stained for E-Cad (red) and MyoII (cyan) undergoing shortening in the Drosophila germband. MyoII is recruited at the junction. Scale bar: 5μm. **E.** Junction stained for E-Cad (red) and MyoII (cyan) undergoing elongation in the Drosophila germband. MyoII is recruited in the anterior and posterior cells, in the vicinity of vertices of newly forming junctions. **F.** Shortening dynamics during a pulse of junctional MyoII. **G.** Elongation dynamics during a pulse of MyoII located in the vicinity of vertices. **H.** Region analyzed to measure MyoII activity during shortening. **I.** Region analyzed to measure MyoII activity during elongation.

The model is in essence timescale-dependent. It is expected that the longer a force is applied, the less reversible the resulting deformation should be. Conversely, the shorter the dissipation timescale is, the faster one will observe irreversible deformations. This is illustrated in Figure 2A, where we show the predicted response of a junction to artificial MyoII pulses in three extreme scenarios. As expected, when the timescale of dissipation τ is much longer than the pulse duration *θ* (that is, when *τ* ≫ *θ*), the response is mainly elastic and deformations are essentially reversible, with complete recovery between pulses (black line). When *τ* ≪ *θ*, the response is mainly viscous, and deformations are essentially irreversible, with no recovery between pulses (light grey line). When *τ* ~ *θ*, the response is viscoelastic, and a partial recovery is observed between pulses (grey line). To test this experimentally, we defined the irreversibility index *I* as the ratio between the irreversible part of the deformation and the maximal deformation (Figure 2B, left panel), and measured *I* as a function of the pulse duration *θ* for N=49 individual pulses (Figure 2B, boxplot). The resulting plot shows that shorter pulses indeed tend to produce more reversible deformations, while longer pulses tend to produce more irreversible deformations, a signature of dissipative mechanics.

**Figure 2.**
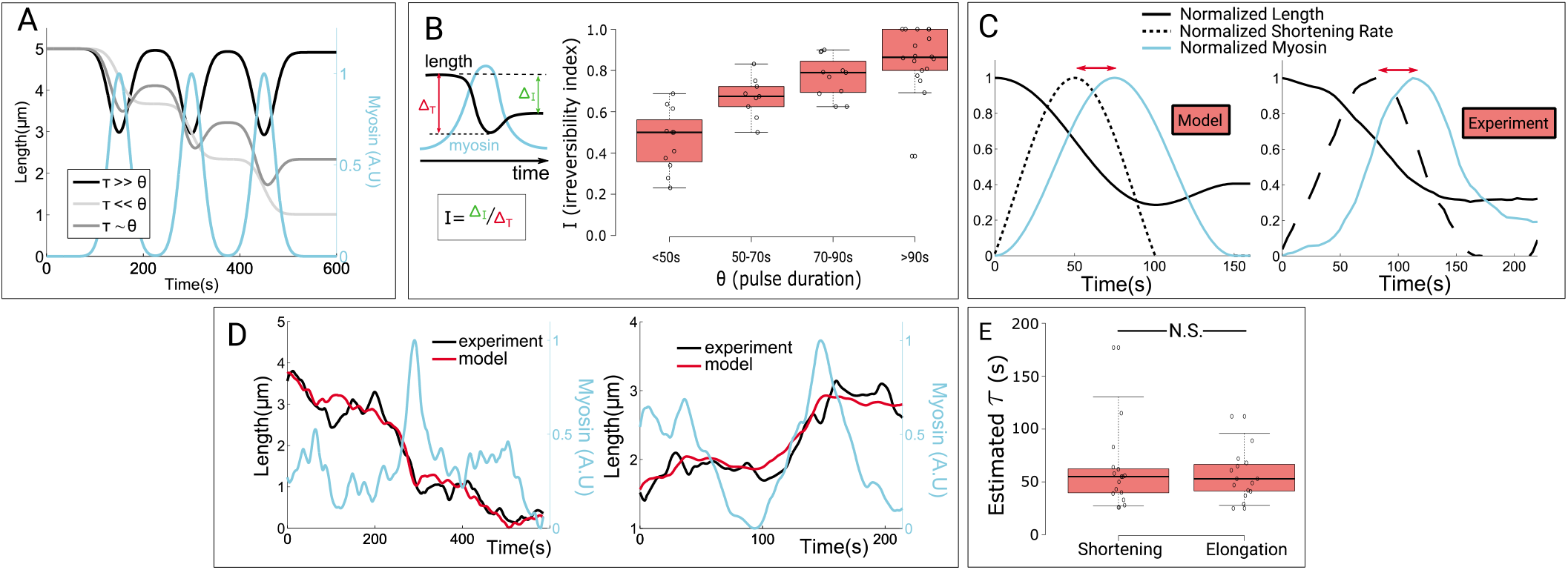
Viscoelastic analysis of contractile pulses. **A.** Shortening dynamics of a model viscoelastic junction in response to artificial Gaussian pulses. Black: *τ* = 1*h* ≫ *θ* (elastic regime), light grey: *τ* = 1*s* ≪ *θ* (viscous regime), grey: *τ* = 1*min* ~ *θ* (intermediate regime). **B.** Left panel: Cartoon of the irreversibility index *I*, defined as the ratio between the irreversible deformation and the maximal deformation. Right panel: Irreversibility index *I* (*θ*) increases with pulse duration *θ*. Center lines of boxes show the medians; boxes limits indicate the 25th and 75th percentiles; whiskers extend 1.5 times the interquartile range from the 25th and 75th percentiles; outliers are represented by dots. **C.** Time delay between the peak of deformation rate (here shortening rate, dotted line) and the peak of MyoII activity (cyan). The former is ahead of the latter, both in a model pulse (left panel) and in experiments (right panel). **D.** Fit of junction shortening (left panel) and junction elongation (right panel), using the experimental MyoII signal as input for the model. **E.** Estimates of τ extracted from fits of experiments, both for shortening (left), and elongation (right). No significant difference is found between shortening and elongation (unpaired t-test, p=0.64). Center lines of boxes show the medians; boxes limits indicate the 25th and 75th percentiles; whiskers extend 1.5 times the interquartile range from the 25th and 75th percentiles; outliers are represented by dots.

**Figure 3.**
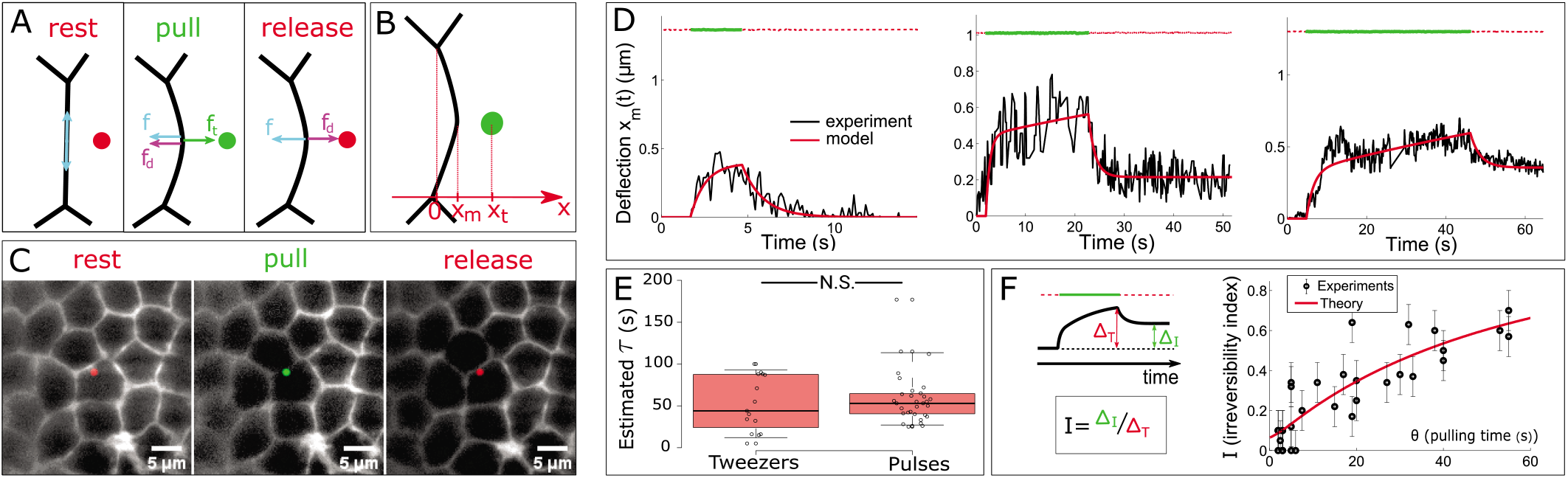
Viscoelastic analysis of optical tweezers experiments. **A.** Cartoon of the pull-release sequence and of the subsequent forces in the optical tweezers experiments. The trap is initially off (red), then switched on (green), and switched off again (red). Forces displayed are the restoring force *f*, the drag force *f_d_*, and the trap force *f_t_*. **B.** Cartoon of deflection quantification. We monitor deflections along the *x* axis, perpendicular to the junction. *x_m_* is the junction position, and *x_t_* the optical trap position. **C.** Timelapse sequence of a pull-release experiment, in which the junction is held for approximately 50s. **D.** Representative kymographs of pull release experiments, with *θ* = 3*s* (left panel), *θ* = 20*s* (middle panel), and *θ* = 40*s* (right panel). The solid black line shows the junction position. The horizontal line displays the optical trap position and its status (on=green, off=dotted red). The solid red line shows the fit obtained from the viscoelastic model. **E.** Estimates of τ extracted from fits of experiments, both for optical tweezers (left), and contractile pulses (shortening and elongation, right). No significant difference is found between tweezers experiments and Myosin pulses (unpaired t-test, p=0.30). Center lines of boxes show the medians; boxes limits indicate the 25th and 75th percentiles; whiskers extend 1.5 times the interquartile range from the 25th and 75th percentiles; outliers are represented by dots. **F.** Left panel: Cartoon of the irreversibility index *I*, defined as the ratio between the irreversible deformation and the maximal deformation. Right panel: Irreversibility index *I* (*θ*) increases with the pulling time *θ*. Black dots show individual experiments. The solid red line shows the analytical prediction.

Two terms contribute to the elongation rate 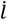 in Eq.1. One scales with MyoII activity *m*, while the other scales with MyoII accumulation rate *ṁ*. Since *m* is pulsatile, maxima of *ṁ* precede maxima of *m*. Hence a straightforward consequence of viscoelasticity is that the maximal rate of elongation 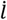 should stand between maxima of *ṁ* and *m*, and therefore precede the maximal MyoII activity (Figure 2C, left panel). We observed that this is indeed the case *in vivo* (Figure 2C, right panel). While it was not interpreted in viscoelastic terms before, this shift was previously reported using temporal cross-correlation analysis, during junction shortening (8), junction elongation (21), and prior to mesoderm invagination when cells’ apical area undergoes MyoII-driven contractions (5). We interpret this consistent shift between MyoII activity and deformation rate as another signature of viscoelastic mechanics.

We have presented semi-quantitative results showing that irreversible deformations generated by MyoII pulses bare the signature of viscoelastic dissipative mechanics. To assess the quantitative accuracy of the model, and hence to estimate the typical dissipation timescale, a direct comparison between experimental and model deformation dynamics is required. To do so, we plugged experimental signals of MyoII activity in our model to predict deformation. We solved Eq.1 for this experimental input and predicted the resulting junction length dynamics *l*(*t*). This prediction depends on the value of model parameters *κ_α_* and *τ*, which we use as adjustable parameters to obtain the best possible fit between the predicted and the experimental *l*(*t*). The best fit is determined by iterating integration of Eq.1 using a gradient descent method. This analysis is performed both for shortening events (Figure 2D, left panel) and elongation events (Figure 2D, right panel). Each fit yields an estimate of the timescale τ. We find a value of 61 ± 9*s* for junction shortening (mean ± standard error), and a similar value of 56 ± 6*s* for junction elongation (Figure 2E). Interestingly, this timescale is slightly shorter than the typical duration of MyoII pulses (typically 80 – 100*s*), which allows dissipation during the pulse, and efficient junction shortening or elongation with little recovery (i.e. reversibility) between consecutive pulses.

Our analysis of MyoII pulses partly relies on the supposed scaling between MyoII activity and force generation. Therefore, we sought to confirm our mechanical analysis using better known forces, applied in a controlled fashion. In a previous study, we introduced optical tweezers as a tool to deform junctions in the embryonic epithelium of *Drosophila*, and to study mechanics on short timescales. Here, we use similar optical forces to perform pull-release experiments on the minute timescale to study the irreversibility of deformations (Figure 3A).

The trap is switched on at a distance *x_t_* of a few hundred nanometers from the midpoint of a junction, then switched off after *θ* seconds. We monitor the deflection *x_m_* (*t*) of the junction from its initial position, that is, *x_m_* (0) = 0 (Figure 3B,C, Movie 1). Assuming a viscoelastic behavior with dissipation on timescale *τ* again yields a Maxwell-like constitutive equation relating the deflection *x_m_*(*t*) to the restoring force *f*(*t*): 
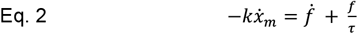
EQN where *k* is the effective elastic stiffness of the junction (see supplementary information for details). Since the trap is instantly switched on or off, the junction undergoes rapid movements and the drag force in the cytosol *f_d_*= −*c_η_ẋ_m_*, where *c_η_* is the drag coefficient, can no longer be neglected. The trap force simply writes *f_t_* = *k_t_*(*x_t_* −*x_m_*), where *k_t_* is the optical trap stiffness (20). Combining this constitutive equation to the force balance *f* + *f_t_* + *f_d_* = 0 yields the following equation of motion:

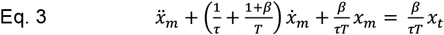
EQN

Note that the inertial term is neglected in Eq.3, since the system is at very low Reynolds number. Eq.3 has three parameters. *β* = *k_t_/k* is dimensionless and compares the trap stiffness to the junction stiffness. It typically controls how far the junction elastically goes towards the optical trap. Note that *β* = 0 when the trap is off; otherwise, our previous study showed that *β* ~ 1 (20). *T* = *c_η_*/*k* is the timescale associated to viscous damping in the cytosol, i.e. the typical time required for the junction to elastically relax into and out of the trap. Our previous study showed that *T* is in the order of one to a few seconds (20). Again, *τ* is the typical dissipation timescale, and underlies the extent of irreversibility.

Consistent with our analysis of contractile pulses, we found that if the duration of force application *θ* is long enough, irreversible deformations are observed. If *θ* is small, typically a few seconds, deformation is mostly reversible (Figure 3D, left panel). For larger values of *θ*, deformations become more and more irreversible (Figure 3D, middle and right panels). In addition, we used the solution to Eq.3 to fit the dynamics of junction deflection in and out of the trap (Figure 3D). Each fit provides an estimate of the three independent parameters *β*, *T*, and *τ*. Consistent with our previous estimates (20), we found that *β* = 1.8 ± 0.7 and *T* = 3 ± 1 *s* (mean ± standard error). More importantly, we obtained *τ* = 50 ± 7 *s*, which is in agreement with the estimates obtained from MyoII pulses analysis (Figure 3E). This suggests that stabilization of the deformation does not require the action of MyoII as a stabilizing force, but is rather a passive consequence of dissipation.

We observed that longer force application times yield deformations that are more irreversible. Again, this was highly expected and in line with a viscoelastic scenario. We defined the rreversibility index *I*(*θ*) as the ratio between the irreversible part of the deflection and the maximal deflection (Figure 3F, left panel). From Eq.3 it is possible to calculate analytically the irreversibility index as a function of the duration of force application *θ* (see supplementary information for details). We measured *I*(*θ*) for *θ* ranging from 2 to 55s (longer times were not experimentally accessible due to cell movements), and compared experimental measurements to the theoretical prediction (Figure 3F, right panel). Note that this prediction has no adjustable parameters, since we directly plugged the values of *β*, *T*, and *τ* obtained earlier in the expression of *I*(*θ*). The good agreement between theory and experiments further confirms our estimate of *τ*, and the relevance of a simple viscoelastic description to interpret junctional mechanics.

## Discussion

The analysis of irreversible cellular deformations resulting from both intrinsic and extrinsic forces reveals that dissipation at cell junctions occurs on the minute timescale in the embryonic ectoderm of *Drosophila*. A direct consequence is that transient morphogenetic forces, such as those produced by pulsatile MyoII activity, can locally generate cell shape changes that are irreversible if they typically last a minute or more. This view is supported by our results showing that longer pulses, or longer tweezing, yield less reversible deformations. Unlike a purely elastic material, junctions submitted to increased forces due to increased MyoII activity dissipate elastic energy. This results in a drift of the reference configuration. Consequently, little or no elastic recoil is observed when MyoII activity drops down to its reference level after a pulse, because pulses typically span on timescales commensurate with the timescale of dissipation. Note that in this viscoelastic scenario, sudden removal of mechanical loading, for instance by laser ablation, would still reveal MyoII-generated stress and yield rapid elastic recoil, as reported before (24,26). Indeed, dissipation of elastic energy does not imply that the system is not prone to mechanical stress, which is continuously generated by MyoII activity. Dissipative mechanics thus provides a straightforward explanation for the so-called mechanical ratchet, suggesting that the stabilization of active junction shortening or elongation might result from passive dissipation without the need of specific stabilizing forces.

We sought to shed light on a series of biological observations with simple physical concepts, rather than to develop a realistic description of the system’s rheology. As stated earlier, a consequence is that the viscoelastic model delineated here is excessively simple, and unlikely to reflect the actual rheological complexity of the system. Indeed, junctions and their cortex are likely to dissipate energy on a distribution of timescales, and possibly involve non-linear viscoelasticity. Yet, our model is sufficient to quantitatively interpret the *in vivo* data accessible to our experiments. More precise rheological measurements would be necessary to develop more sophisticated rheological models. This remains experimentally very challenging, especially *in vivo*.

Large deformations at the scale of the tissue are likely to involve a variety of additional mechanical contributors, possibly including the surrounding yolk and vitelline membrane, as well as topological transitions between cells. These are not considered in this study, which is limited to the junction scale. The formation of new antero-posterior junctions, because they modify tissue topology, might for instance be required for long-term stabilization of large, tissue-scale elongation, once MyoII polarity is lost. Such contributions, emerging at the scale of several cells, affect the mechanics of the system and thereby are likely to involve effective dissipation timescales at the scale of the tissue. Note that in suspended monolayers, which display no or very few topological transitions and have no substrate, stress relaxation at the tissue scale typically occurs on the minute timescale (19), consistent with our estimate. Although this is out of the scope of this study, continuum mechanics models might provide a more adequate framework to describe the full tissue dynamics (13,27). Bridging the gap between subcellular analysis such as ours, and macroscopic models of tissue flows, which can incorporate effective mechanical parameters as well as a wide variety of cell behaviors (topological transitions, growth, division, apoptosis, etc.), remains a challenge for the physics of tissue morphogenesis.

The question as to how elastic energy is dissipated at the molecular level is not addressed here, and remains rather open. Turnover of cortical components, by remodeling the stress-bearing structure, is likely to allow cells to adapt to a new geometry or mechanical environment. Therefore the turnover rate of Actin and Actin crosslinkers has been proposed to underlie dissipation (11,12,28). MyoII action has also been shown to fluidize the cytoskeletal network (29,30) and enhance Actin disassembly (31). The redistribution of adhesion molecules at deformed junctions might also be important to stabilize deformations, and Cadherin rate of endocytosis might thus be a limiting factor for stabilization. Interestingly, such contributions of protein dynamics to dissipation allow cell-, tissue-or organ-specific regulation of the dissipation timescales. This offers a versatile tool for morphogenesis: besides the ability of cells to generate localized and/or polarized forces, local and/or polarized control of the mechanical response to forces become possible, not only in a quantitative manner (stiffer or softer), but in a qualitative manner (more elastic or more viscous). Tissue-specific tuning of dissipative properties might also be important to ensure that short-lived or abnormal mechanical stresses do not produce dramatic irreversible deformations. In the system investigated here, the timescale of force generation, that is, the timescale of MyoII pulses, is commensurate with the typical dissipation timescale, which seems crucial to achieve efficient deformations. This observation might be relevant to other morphogenetic systems, although further work is required to confirm this hypothesis. In particular, other morphogenetic events displaying reversible and irreversible contractions should be analyzed. For example, actomyosin pulses during zebrafish optic cup morphogenesis drive essentially reversible shortening events of cells’ apico-basal axis length (32). Interestingly and in contrast with cells investigated here, shortening and Myosin oscillations seem synchronous, which is consistent with a reversible, elastic regime of deformation. Similarly, during the first phase of dorsal closure, pulses of MyoII in the medio-apical cortex cause reversible contractions of amnioserosa cells’ apical area (33,34). This is followed by a second phase during which pulses become more efficient, with net shrinkage after each pulse. A possible mechanism is a gradual increase of dissipation during the process, causing pulses to first probe an elastic regime, while as time passes they probe an increasingly viscous regime, allowing more efficient deformations.

Altogether, our results provide a first *in vivo* estimate of the typical dissipation timescale associated with junctional remodeling, and shed a new light on the mechanism of the so-called mechanical ratchet, highlighting the key contribution of dissipation in stabilizing deformations. It opens a new avenue for the understanding of tissue morphogenesis and the irreversibility of deformations by an emphasis on viscoelastic, dissipative properties of cells, beyond force generation per se.

## Acknowledgements

We thank Claire Chardès for assistance with the optical tweezers set up, and members of the Lenne and Lecuit groups for stimulating and useful discussions during the course of this project.

This work was supported by an FRM Equipe Grant FRM DEQ20130326509, Agence Nationale de la Recherche ANR-Blanc Grant, Morfor ANR-11-BSV5-0008 (to P.-F.L.), the Labex INFORM (ANR-11-LABX-0054) of the A*MIDEX project (ANR-11-IDEX-0001-2), funded by the ‘Investissements d’Avenir French Government program’. We acknowledge France-BioImaging infrastructure supported by the French National Research Agency (ANR–10–INSB-04-01, «Investments for the future»).

## Author contributions

RC and PFL conceived the project, and discussed it with CC and TL. CC and BD performed Myosin pulses experiments; and CC, BD and RC analyzed the data. RC performed the optical tweezers experiments and analyzed the data. RC designed the model. All authors discussed the data. RC wrote the manuscript and all authors commented on it.

